# Species-specific barriers constrict Orsay virus host range across the *Caenorhabditis* genus

**DOI:** 10.64898/2026.04.10.717618

**Authors:** Dominik Herek, Juan C. Muñoz-Sánchez, Ana Villena-Giménez, Victoria G. Castiglioni, Santiago F. Elena

**Author notes:** To whom correspondence should be addressed: Emails: V.G.C., S.F.E.

## Abstract

Predicting the host range and spillover potential of RNA viruses requires understanding how ecological, immunological, and evolutionary factors jointly shape viral life cycles across related hosts. Here we integrate population-level viral load dynamics, single-animal heterogeneity, tissue-level progression, transmission competence, and evolutionary sustainability to map the eco evolutionary barriers that Orsay virus encounters across six *Caenorhabditis* species. We show that host species identity determines the timing and completeness of the viral life cycle, producing species specific combinations of susceptibility, replication kinetics, RNA2 to RNA1 stoichiometric balance, virion egress, and onward transmission. These phenotypes correspond to distinct host competence phenotypes, ranging from permissive (*Caenorhabditis elegans*) to restrictive or evolutionarily dead end host species. Alternative host species disrupt viral life cycle synchrony through delayed replication, truncated cycles, or failure to produce lumen localized virus, thereby reducing transmission and preventing viral adaptation upon serial passage. Our results demonstrate how temporal mismatches between viral replication and host physiology create a series of eco evolutionary barriers to emergence, offering a mechanistic framework for predicting viral host range.

## Introduction

Emerging infectious diseases typically arise when pathogens expand their host range, crossing ecological, physiological, and evolutionary boundaries (Plowright et al. 2017; Holmes 2022). Spillover succeeds only when a pathogen sequentially clears a series of barriers, from exposure and cellular entry to intracellular replication, immune evasion, within-host amplification, shedding, and onward transmission (Plowright et al. 2017; Borremans et al. 2019). An eco-evolutionary perspective emphasizes that establishment in a new host depends not merely on molecular compatibility but on the alignment of viral life-cycle kinetics with host developmental, immunological and ecological traits (Longdon et al. 2011). When timing, genomic and proteomic stoichiometries, or tissue tropism are out of synchronization, infections stall or remain abortive, preventing transmission.

Pandemics underscore the stakes of getting this right (Short et al. 2018; Mallah et al. 2021). Many emerge from zoonotic jumps, especially by RNA viruses, yet successful adaptation and sustained spread after cross-species contact are rare because host defenses impose formidable barriers; spillover therefore occurs more often between closely related hosts (Manzin et al. 2000; van Heuverswyn and Peeters 2007; Jones et al. 2008; Woolhouse et al. 2013; Plowright et al. 2017; Wasik et al. 2019; Andersen et al. 2020; McKeown et al. 2025). RNA viruses are nonetheless well positioned to overcome these obstacles given their short generation times, large population sizes, and high mutation rates that bestow them great evolvability and accelerate exploration of adaptive space (Sanjuán et al. 2010; Elena et al. 2011; Holmes 2022). Understanding viral dynamics across related hosts is therefore essential to anticipate host range and spillover potential.

Most prior work is retrospective or diversity surveys that are invaluable for surveillance yet limited for hypothesis testing (Luis et al. 2013; Meadows et al. 2023). Controlled experimental models offer a complementary route: phage-bacteria and cell culture have yielded key insights but lack whole-organism ecological context (Clarke et al. 1993; Turner and Elena 2000; Duffy et al. 2007; Nguyen et al. 2012; Shukla et al. 2012; Morley et al. 2015; Iketani et al. 2018). This limitation has been successfully overcome by the use of plant viruses (Hillung et al. 2014; Willemsen et al. 2017; González et al. 2019), although these models still lack close relevance to animal hosts. Mammalian models can directly link viral load to transmission but are ethically and logistically demanding, limiting scale (Kim et al. 2015; Bashor et al. 2021; Pulit-Penaloza et al. 2023). Together, these gaps motivate tractable, organismal systems that can measure spillover-relevant traits (*e.g*., replication timing, tissue tropism, immune clearance, shedding, and transmission) across closely related hosts under standardized conditions.

The *Caenorhabditis* - Orsay virus (OrV) pathosystem is a tractable, whole-organism model for mechanistic dissection of spillover barriers across closely related hosts. *Caenorhabditis elegans* is a free-living bacterivore commonly found on rotting plant matter and is easy to culture with powerful genetic tools (Félix and Braendle 2010). It is a mainstay for studies of development (Hubbard and Greenstein 2000), aging (Shen et al. 2018) and innate immunity (Ermolaeva and Schumacher 2014). The identification of OrV as a natural pathogen of *C. elegans* (Félix et al. 2011) opened the door to *in vivo* analyses of host-virus interactions, antiviral innate responses and virus experimental evolution (Sowa et al. 2020; Batachari et al. 2024; Castiglioni et al. 2024a, 2024b, 2025). This blend of ecological relevance, genetic tractability, and comparative breadth across the genus makes *Caenorhabditis* an ideal platform to test eco-evolutionary predictions about host range and spillover (Shaw and Kennedy 2022, 2025).

OrV is currently the only known virus that naturally infects *C. elegans*. It is a positive-sense, single-stranded RNA virus with two genomic segments. RNA1 encodes the RNA-dependent RNA polymerase (RdRp), and RNA2 encodes the capsid protein (CP), the δ protein required for nonlytic egress, and the CP-δ fusion involved in viral entry (Félix et al. 2011; Jiang et al. 2014a; Yuan et al. 2018; Guo et al. 2020). Infections are typically restricted to one or few intestinal cell (Castiglioni et al. 2024a). *C. elegans* counters OrV via RNA interference, RNA uridylation, and the intracellular pathogen response (IPR) (Ashe et al. 2013; Reddy et al. 2017; Le Pen et al. 2018). Fitness costs in *C. elegans* are modest, a slight reduction in fertility and lifespan (Ashe et al. 2013; Castiglioni et al. 2025). Across mutants and environments, OrV consistently shows an early replication phase with a viral load peak around 12 h post-inoculation (hpi), whereas later-stage dynamics (peak magnitude, infection rate, number of infected cells, and RNA1-only *vs* RNA1 + RNA2 states) vary widely (Castiglioni et al. 2024a, 2025; Melero et al. 2025; Villena-Giménez et al. 2026). Together, these patterns indicate that both viral and host factors shape infection outcomes, motivating a cross-species, eco-evolutionary analysis to pinpoint the barriers that limit OrV emergence in alternative *Caenorhabditis* hosts.

The *Caenorhabditis* genus comprises > 65 species with substantial genetic diversity (Kiontke et al. 2004; Stevens et al. 2020). The discovery that several are OrV-susceptible established the *Caenorhabditis* - OrV system as a tractable model for spillover. Comparative work shows that susceptibility varies widely across *Caenorhabditis*, with species and strains ranging from permissive to strongly resistant (Félix and Wang 2019; Shaw and Kennedy 2022; Alkan et al. 2024). Yet we still lack a mechanistic, multiscale picture of how the OrV life cycle proceeds, or fails, across hosts. Prior work emphasized predicting establishment by modeling infection intensity, prevalence and shedding (Shaw and Kennedy 2025); our study complements this by resolving acute infection events and cross-species transmission dynamics, the critical pieces needed for a causal, eco-evolutionary account of viral emergence. Bridging this gap requires integrating population dynamics, cell-level progression, within-host heterogeneity, and evolutionary sustainability to connect viral phenotypes with eco-evolutionary theory of host range.

Here we apply a unified experimental framework to six *Caenorhabditis* species, combining quantitative viral kinetics, single-animal distributions, smiFISH-based tissue localization, transmission assays, virulence measurements, and experimental evolution. We find that host species imposes distinct temporal and mechanistic constraints on the viral life cycle, shaping susceptibility, replication efficiency, RNA-segment stoichiometry, egress, transmission probability, and the capacity for adaptation. Placed within a spillover-barrier framework, these patterns resolve host-competence phenotypes and show how life-cycle mismatches generate eco-evolutionary barriers to emergence. This approach builds on theory that frames host range as the outcome of hierarchical barriers and trait mismatches, modulated by phylogenetic proximity, contact structure, and viral traits that broaden host breadth (Geoghegan et al. 2016; Olival et al. 2017; Plowright et al. 2017; Becker et al. 2019). It also incorporates individual-level heterogeneity (*e.g*., superspreading) and community-level competence, which can decouple prevalence from transmission (Lloyd-Smith et al. 2005; Kilpatrick et al. 2006), and recognizes that landscape and movement shape outbreak thresholds while clade-specific life histories influence zoonotic potential (White et al. 2018; Letko et al. 2020).

## Results

We characterize OrV infection across five alternative hosts: *Caenorhabditis tropicalis* JU1428, *Caenorhabditis wallacei* JU1873, *Caenorhabditis remanei* SP8, *Caenorhabditis macrosperma* JU1857, and *Caenorhabditis sulstoni* SB454. *C. elegans*, *C. tropicalis*, *C. wallacei*, and *C. remanei* belong to the *Elegans* group, while *C. macrosperma* and *C. sulstoni* pertain to the *Japonica* group (Stevens et al. 2020).

### Barrier 1: Susceptibility and entry

To examine host susceptibility in the selected species (Shaw and Kennedy 2022), we performed smiFISH. Synchronized animals inoculated at hatching were collected at 24 hpi for all species and additionally at 14 hpi for *C. elegans* and *C. remanei*, and the proportion of animals with a positive smiFISH signal was determined. Infection was detected at 24 hpi in every host species, except *C. remanei*, in which infection was observed only at 14 hpi (Fig. 1A - B), consistent with the viral accumulation dynamics shown in Fig. 2E (see Barrier 2 below). Host species had a significant effect on the infection rate (Fig. 1C; 24 hpi: χ^2^ = 308.571, 4 d.f., *P* < 0.0001; 14 hpi: χ^2^ = 223.319, 1 d.f., *P* < 0.0001). At 24 hpi, *C. elegans* showed the highest proportion of infected animals (64.0%), whereas all other species exhibited significantly lower infection rates: *C. tropicalis* 6.3%, *C. wallacei* 1.8%, *C. macrosperma* 4.9%, and *C. sulstoni* 9.3% (*P* < 0.0001 for all comparisons). At 14 hpi, infection was detected in only 5.2% of *C. remanei* animals, again significantly lower than in *C. elegans* (79.5%, *P* < 0.0001).

**Figure 1.**
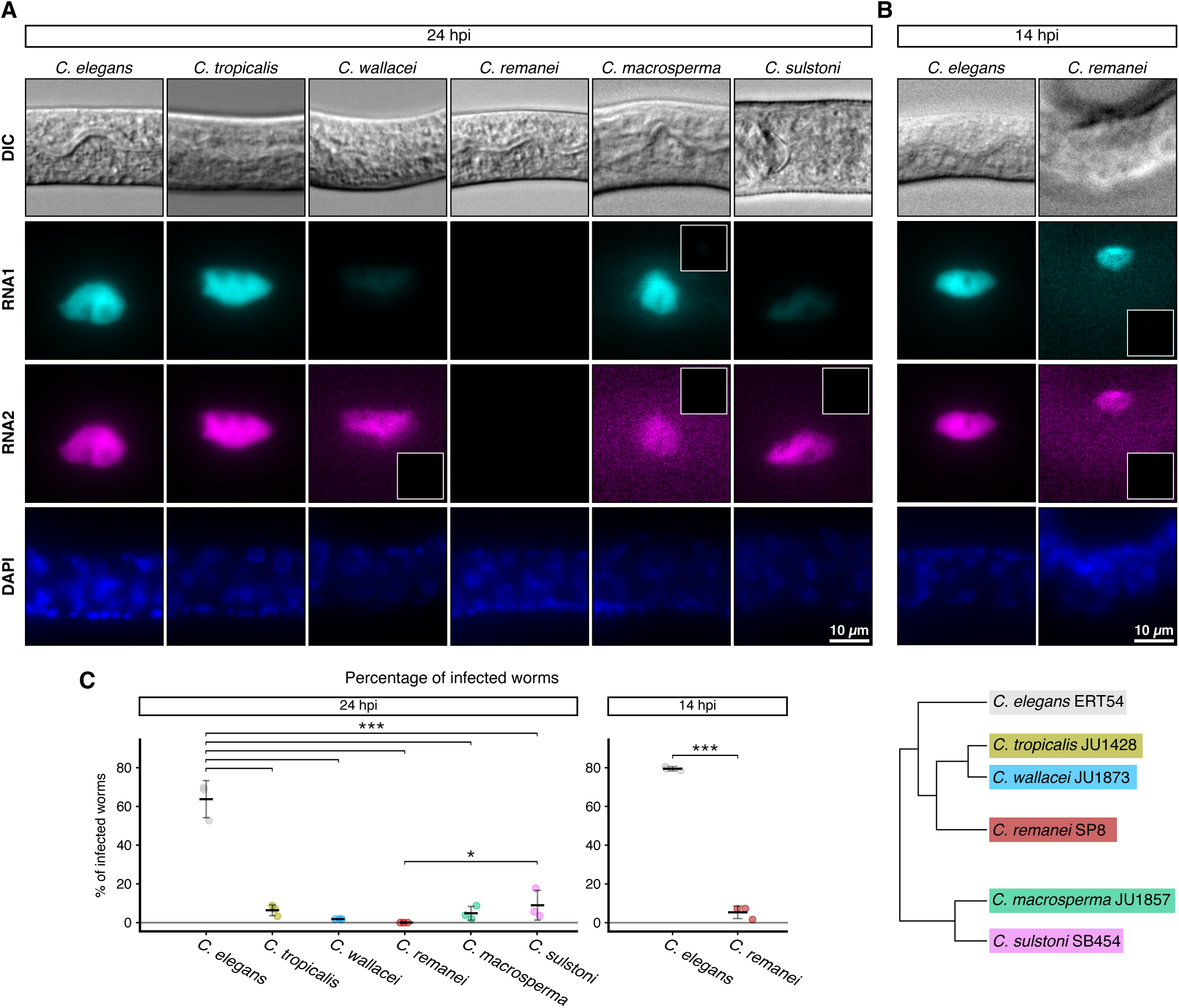
Representative smiFISH images of OrV infection in alternative host species. (**A**) *C. elegans* and the alternative host species at 24 hours post inoculation (hpi). (**B**) *C. elegans* and *C. remanei* at 14 hpi. The brightness levels of all images were equalized to allow for comparisons. All images are displayed with the same brightness and contrast levels to allow for visual comparisons, except for the images that have insets. The inset displays same brightness and contrast as the other images, and the magnified images have increased brightness to visualize low signal. Infection was not detected in any *C. remanei* worms at 24 hpi. Top: differential interference contrast (DIC), OrV RNA1 (cyan), OrV RNA2 (magenta), DAPI (blue). (**C**) Percentage of infected worms at 24 and 14 hpi determined by counting the number of animals with a positive smiFISH signal (*n* = 3 replicates of 56 ±4 animals). The legend showing colors used for each species also shows phylogenetic relationships adapted from Stevens et al. (2020).

**Figure 2.**
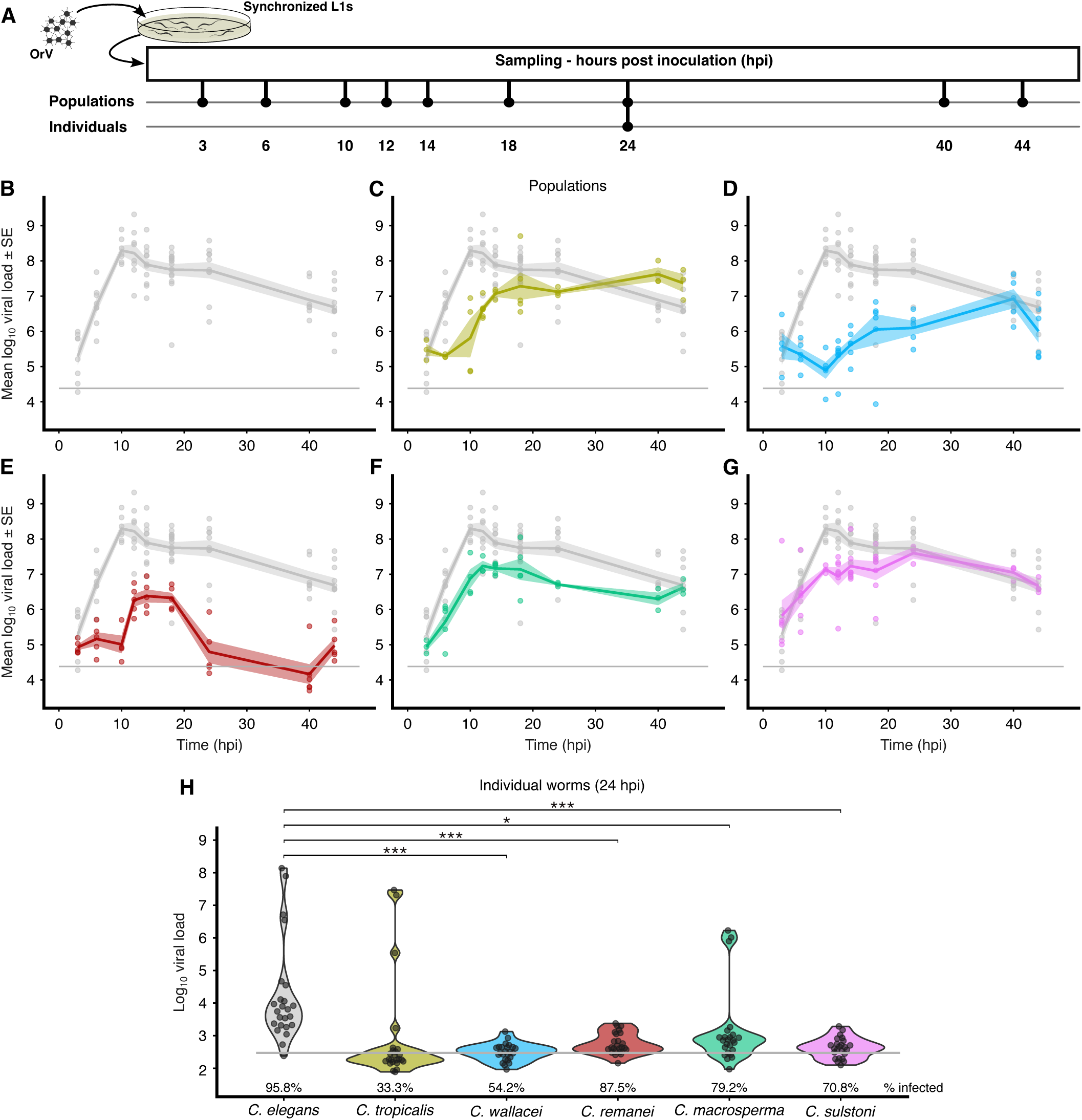
OrV viral load dynamics and single worm viral load distributions across host species. (**A**) Schematic explanation of the experimental design. Synchronized populations of ∼500 worms were inoculated at hatching with 2.8×10^9^ copies of OrV RNA2 and sampled at the indicated timepoints for total RNA extraction or single worm lysis and OrV viral load by measured by absolute RT-qPCR. Viral load over time throughout the first 44 hpi in the different host species: (**B**) *C. elegans* ERT54 (grey), (**C**) *C. tropicalis* (yellow), (**D**) *C. wallacei* (blue), (**E**) *C. remanei* (red), (**F**) *C. macrosperma* (green), and (**G**) *C. sulstoni* (pink). Viral load was measured as the number of OrV RNA2 copies in 10 ng of total RNA; 3 - 16 replicates per strain per timepoint. (**H**) Violin plots of OrV viral load of individual animals (*n* = 24 for each species) at 24 hpi. Viral load was measured as the number of OrV RNA2 copies in 1/5 of the single worm lysate. Horizontal grey lines represent the technical detection limit (mean + 2SD of negative controls).

In addition to the frequency of detectable infection, early failures in the replication program can signal entry/establishment bottlenecks. At 14 hpi, we observed substantial fractions of cells with RNA1 only (see Barrier 3 below), consistent with abortive early infection and/or rapid host clearance in specific species (notably *C. remanei*), further supporting the presence of strong early barriers to successful entry and initiation of the full viral cycle.

### Barrier 2: Within-host replication dynamics

To better understand the replication dynamics of OrV in the alternative hosts, we measured viral load during the first 44 hpi. For this purpose, synchronized populations of ∼500 animals were inoculated at hatching and sampled at nine time points (Fig. 2A). Viral load was quantified as the number of OrV RNA2 copies per ng of total RNA using the absolute RT-qPCR method. The viral load trajectory observed in *C. elegans* (Fig. 2B) matched our previous results (Castiglioni et al. 2024a). Among the different host species, viral load dynamics differed significantly, as did the interaction between host species and time (Fig. 2B - G; gray horizontal line indicates the technical detection cutoff; χ^2^ = 281.081, 4 d.f., *P* < 0.0001; χ^2^ = 302.444, 48 d.f., *P* < 0.0001, respectively).

Averaged across all time points, *C. elegans* exhibited the highest viral load of all host species (*C. tropicalis*, −77.4%, *P* < 0.0001; *C. wallacei*, −97.0%, *P* < 0.0001; *C. remanei*, −98.9%, *P* < 0.0001; *C. macrosperma*, −82.7%, *P* < 0.0001; and *C. sulstoni*, −59.6%, *P* = 0.0008). Moreover, the alternative hosts showed differences in the duration of the eclipse phase of viral accumulation compared to the natural host. Slopes estimated from an exponential growth model using the first 10 hpi were lower for all alternative hosts relative to *C. elegans* (*C. tropicalis*, −86.9%, *P* = 0.0001; *C. wallacei*, −123.3%, *P* < 0.0001; *C. remanei*, −97.4%, *P* < 0.0001; *C. macrosperma*, −34.5%, *P* = 0.0317; and *C. sulstoni*, −56.0%, *P* = 0.0089). These results suggest a possible desynchronization of the viral replication cycle, with a prolonged lag phase and/or slower exponential growth rate in the alternative hosts. Viral load remained high throughout the entire 44 hpi observation period in all species except *C. remanei*, which showed only a short replication period followed by viral clearance at 24 hpi. Overall, these results indicate that OrV replication occurs with species-specific dynamics that are slower, reach lower peaks, and reflect a temporal shift in the viral cycle relative to the natural host, *C. elegans*.

To further examine inter-individual differences in the ability of *C. elegans* and alternative hosts to support viral replication and accumulation, we inoculated synchronized animals at hatching and measured the viral load of individual animals at 24 hpi. After discarding individuals with values below the cutoff threshold (mean + 2 SD of RT-qPCR values from noninfected animals), we found significant differences in viral load among host species (Fig. 2H; χ^2^ = 30.757, 4 d.f., *P* < 0.0001). *C. elegans* and *C. tropicalis* formed a homogeneous group with higher viral load than the remaining four species (*C. wallacei*, −97.7%, *P* = 0.0001; *C. remanei*, −96.7%, *P* = 0.0001; *C. macrosperma*, −88.5%, *P* = 0.0389; *C. sulstoni*, −97.1%, *P* = 0.0001).

Viral load among individuals of the same species showed significantly right-skewed distributions in all cases: *C. elegans* (range: 5.3×10^2^ - 1.4×10^8^; *g*□ = 3.537 ±0.481, *P* < 0.0001), *C. tropicalis* (3.2×10^2^ - 2.9×10^7^; *g*□ = 1.611 ±0.752, *P* = 0.0322), *C. wallacei* (3.0×10^2^ - 1.3×10^3^; *g*□ = 2.368 ±0.616, *P* = 0.0001), *C. remanei* (3.0×10^2^ - 2.4×10^3^; *g* = 1.247 ±0.501, *P* = 0.0129), *C. macrosperma* (3.0×10^2^ - 1.69×10^6^; *g*□ = 2.552 ±0.524, *P* < 0.0001), and *C. sulstoni* (3.1×10^2^ - 1.9×10^3^; *g*□ = 1.857 ±0.550, *P* = 0.0007). To test whether bimodal Lognormal models better fit our data than a unimodal single Lognormal distribution (Down et al. 2020), we compared fits between the two approaches (Table 1, Fig. S1A - F). Based on Akaike weights (> 0.95), viral load data from *C. wallacei* and *C. sulstoni* were well explained by single Lognormal distributions (Fig. S1C, F). In contrast, model fit improved significantly for *C. elegans*, *C. tropicalis* and *C. macrosperma* when using a mixture of two Lognormals (Fig. S1A, B, E), despite the addition of three parameters (Table 1). Statistical power was not enough to decide among models in the case of *C. remanei* (Fig. S1D). For *C. elegans*, 83.3% of individuals fell into a distribution with a mean of 4.0×10^3^, whereas the remaining high producers followed a distribution with a mean of 2.1×10^7^. For *C. tropicalis*, 87.5% belonged to a distribution with a mean of 2.1×10^2^, and high producers followed a distribution with a mean of 5.9×10^6^. For *C. macrosperma*, 87.5% produced low viral loads (mean 5.2×10^2^), while high producers followed a distribution with a mean of 1.1×10^6^. These results suggest that the abundance of high producers varies across species, having a direct consequence on the differences in the prevalence of infections among host species.

**Table 1.**
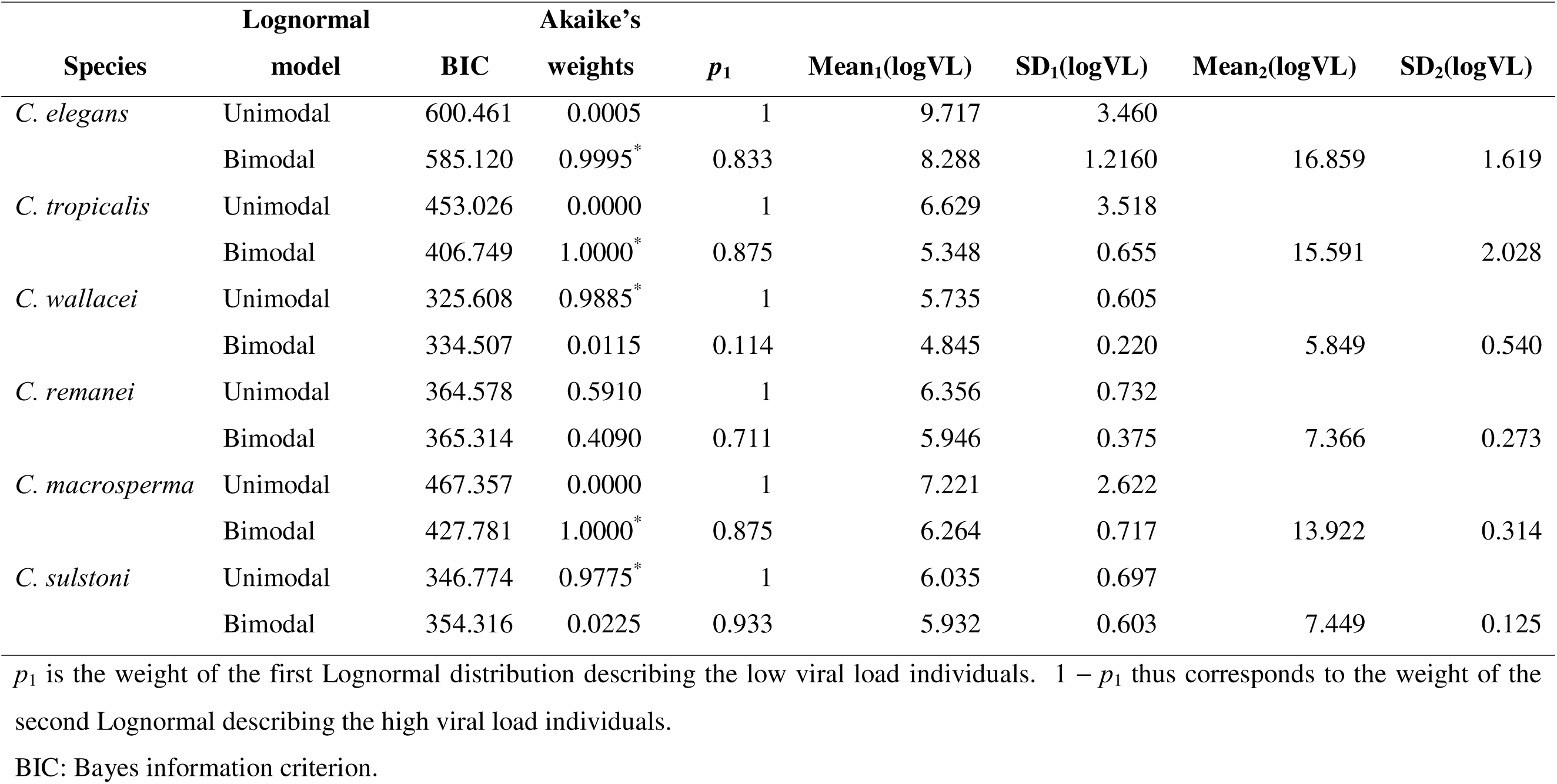
Fitting of viral loads (VL) to unimodal or bimodal Lognormal models.

### Barrier 3: Viral genome stoichiometry and life-cycle progression

To clarify the stage of the viral cycle in each host species, we further analyzed our previously obtained smiFISH dataset. We first examined whether infected cells contained only RNA1 or both RNA segments. At both time points, most infected animals and all animals with virus in the lumen carried both RNAs. At 14 hpi, however, significant differences emerged in the number of animals lacking RNA2 between *C. elegans* (4/138) and *C. remanei* (4/9) (Fig. 3A; Fisher’s exact test, *P* = 0.0004). The very low odds ratio (0.050) indicates a strong bias toward *C. remanei* individuals carrying only RNA1, providing additional evidence of abortive OrV infection in this species. By 24 hpi, the number of animals missing RNA2 was lower (*C. elegans* 2/123, *C. tropicalis* 1/9, *C. wallacei* 0/3, *C. macrosperma* 0/8, and *C. sulstoni* 1/16) and consistent across species (Fisher’s exact test, *P* = 0.2634). This pattern indicates that infection with both RNA segments is more likely to avoid clearance by the host during early time points.

**Figure 3.**
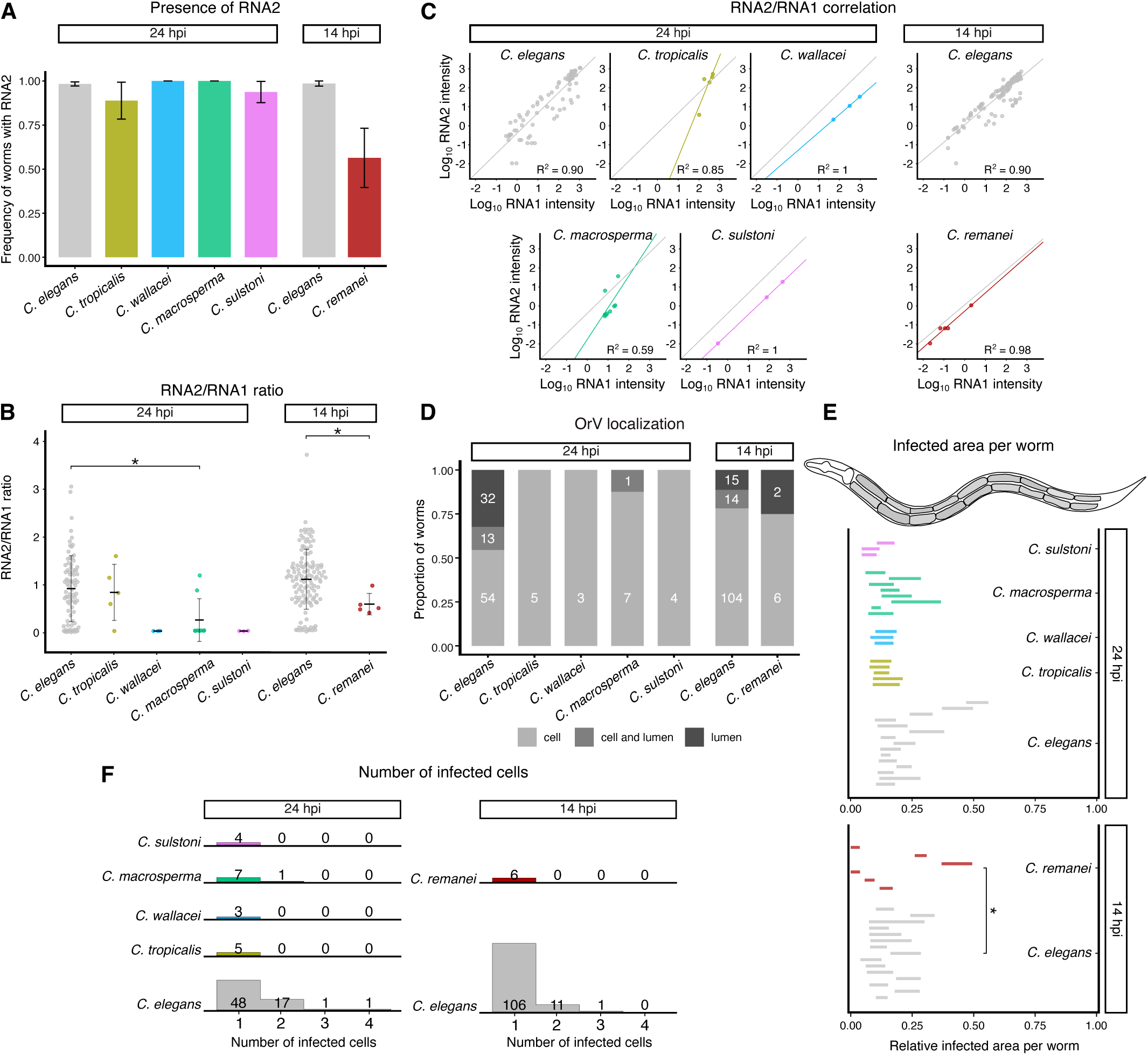
Spatial localization and RNA ratios of OrV infection in alternative host species. (**A**) Bar plot of the proportion of animals in which OrV RNA2 signal was detected. Error bars represent the standard error of the Binomial proportion (*C. elegans*, grey; *C. tropicalis*, yellow; *C. wallacei*, blue; *C. remanei*, red; *C. macrosperma*, green; *C. sulstoni*, pink). (**B**) OrV RNA2:RNA1 ratios at 24 and 14 hpi. OrV RNA2:RNA1 ratios were calculated based on the normalized median intensity values of RNA1 and RNA2 smiFISH signal. (**C**) Correlations plots of RNA1 and RNA2 intensities in each species and timepoint. As a reference, the regression line of the corresponding *C. elegans* timepoint is shown as a grey line in all panels. Intensities are displayed in arbitrary units. (**D**) OrV localization within cells, intestinal lumen, or cell and lumen at 24 and 14 hpi. Bar plot shows the proportion of infected worms in each category within each species and timepoint. The numbers inside the bar plot denote the number of worms in each category. (**E**) Infected area and position of infection per animal relative to intestine length at 24 and 14 hpi. At 14 hpi, *C. elegans* and *C. remanei* differ significantly in the size of the relative infected area (10.4% *vs* 5.7%, *P* = 0.0014). (**F**) Number of infected cells per animal at 24 and 14 hpi. The histograms represent the relative abundance of each category within each species and timepoint.

We previously showed that the RNA2:RNA1 ratio changes dynamically during infection in *C. elegans* (Castiglioni et al. 2024a). As an additional measure of viral cycle progression, we quantified the RNA2:RNA1 ratio in each species by comparing their respective normalized smiFISH fluorescence intensities. RNA2:RNA1 ratio was significantly higher in *C. elegans* (1.1 ±0.6) than in *C. remanei* (0.6 ±0.2) at 14 hpi (Fig. 3B; 46.3%, *P* = 0.0228). At 24 hpi, overall significant differences in the ratio were also found (Fig. 3B; χ² = 16.423, 4 d.f., *P* = 0.0025) among species, with *C. elegans* still showing the highest mean ratio (0.92 ±0.07). However, pairwise differences between *C. elegans* and *C. macrosperma* (0.3 ±0.2) showed the only significant case (*P* = 0.0345). (In all other comparisons: *C. tropicalis* 0.8 ±0.3, *P* = 1; *C. wallacei* 0.04 ±0.37, *P* = 0.1652; and *C. sulstoni* 0.04 ±0.37, *P* = 0.0512.) The measured RNA1 and RNA2 intensities were strongly correlated after controlling for host species and time post-inoculation (Fig. 3C; *r* U = 0.787, 232 d.f., *P* < 0.0001). Most of the cells of the alternate species fell below the *C. elegans* regression line (shown in grey), indicating that the ratio of RNA2:RNA1 in these hosts is skewed towards a greater amount of RNA1 compared to the *C. elegans* average. Only a few cells in *C. macrosperma* and *C. tropicalis*, the two species with high-viral load individuals, fell above the *C. elegans* line; however, all observed values were within the variability of observations in *C. elegans*. These results indicate that, depending on the host species, OrV is less efficient at replicating RNA2 relative to RNA1 and therefore may be impaired in producing new virions.

### Barrier 4: Virion egress and tissue localization

To further test our hypothesis of desynchronized viral cycles, we examined the localization of OrV, classifying infection as viral RNAs being detected within cells, in the intestinal lumen, or in both (Fig. S2A). Presence of virus in the lumen indicates successful completion of the viral cycle and the potential for onward transmission. We found significant differences only at 24 hpi: *C. elegans* showed a much higher proportion of animals with virus in the lumen or in both compartments (lumen: 32/99; cell + lumen: 13/99), whereas the other species showed almost exclusively cell-restricted infection (Fig. 3D; χ^2^ = 10.073, 3 d.f., *P* = 0.0180). The only alternative host in which lumen-localized virus was observed was *C. macrosperma* (cell + lumen: 1/8). As expected, at 14 hpi, *C. elegans* exhibited a significantly lower proportion of lumen-localized virus (lumen: 15/133; cell + lumen: 14/133) compared with 24 hpi (Fig. 3D; χ^2^ = 16.552, 1 d.f., *P* < 0.0001). A small number of *C. remanei* individuals also displayed lumen-localized virus at 14 hpi (2/6). Taken together with our transmission results, the higher frequency of luminal virus observed in *C. elegans* and *C. remanei* indicate a temporal shift in the viral life cycle which delays or prevent egression.

To complement localization, we quantified infected area and position along the intestine and number of infected cells per animal. At 24 hpi, no significant differences were detected among species in relative infected area (Fig. 3E; ∼10.1% of the measured area infected, χ^2^ = 3.079, 3 d.f., *P* = 0.3796), and no significant positional differences were observed either (χ^2^ = 7.502, 3 d.f., *P* = 0.0575). At 14 hpi, the infected area in *C. remanei* (5.7%) was smaller than in *C. elegans* (10.4%) (*P* = 0.0143), although no significant positional differences were observed (*P* = 0.6222). The number of infected cells did not differ significantly among species at 24 hpi (Fig. 3F; χ^2^ = 0.680, 3 d.f., *P* = 0.8780) or between *C. elegans* and *C. remanei* at 14 hpi (Fig. 3F; χ^2^ = 0.065, 1 d.f., *P* = 0.7992). However, individuals containing two infected cells were most common, and animals with three or four infected cells were observed only in *C. elegans*, not in any alternative hosts (Fig. S2B), mirroring the higher viral load outliers observed more frequently in *C. elegans* (Barrier 2).

### Barrier 5: Transmission competence

To investigate how differences in OrV replication and viral cycle dynamics affect epidemiological parameters in alternative host species, we measured the probability of OrV transmission from each host species to *C. elegans*. To do this, we transferred 20 inoculated, synchronized animals at 14 or 24 hpi into populations of 111 ±26 healthy *C. elegans* ERT54 animals (Fig. 4A). These two time points were selected because they correspond to the initial release of new OrV virions (14 hpi) and a more advanced stage of the viral cycle near clearance (24 hpi) in *C. elegans* (Castiglioni et al. 2024a). The ERT54 strain contains an integrated fluorescent infection reporter that enables real-time tracking of infections by fluorescence microscopy (Bakowski et al. 2014). Twenty-four hours after transfer, we quantified the percentage of ERT54 animals expressing the infection reporter.

**Figure 4.**
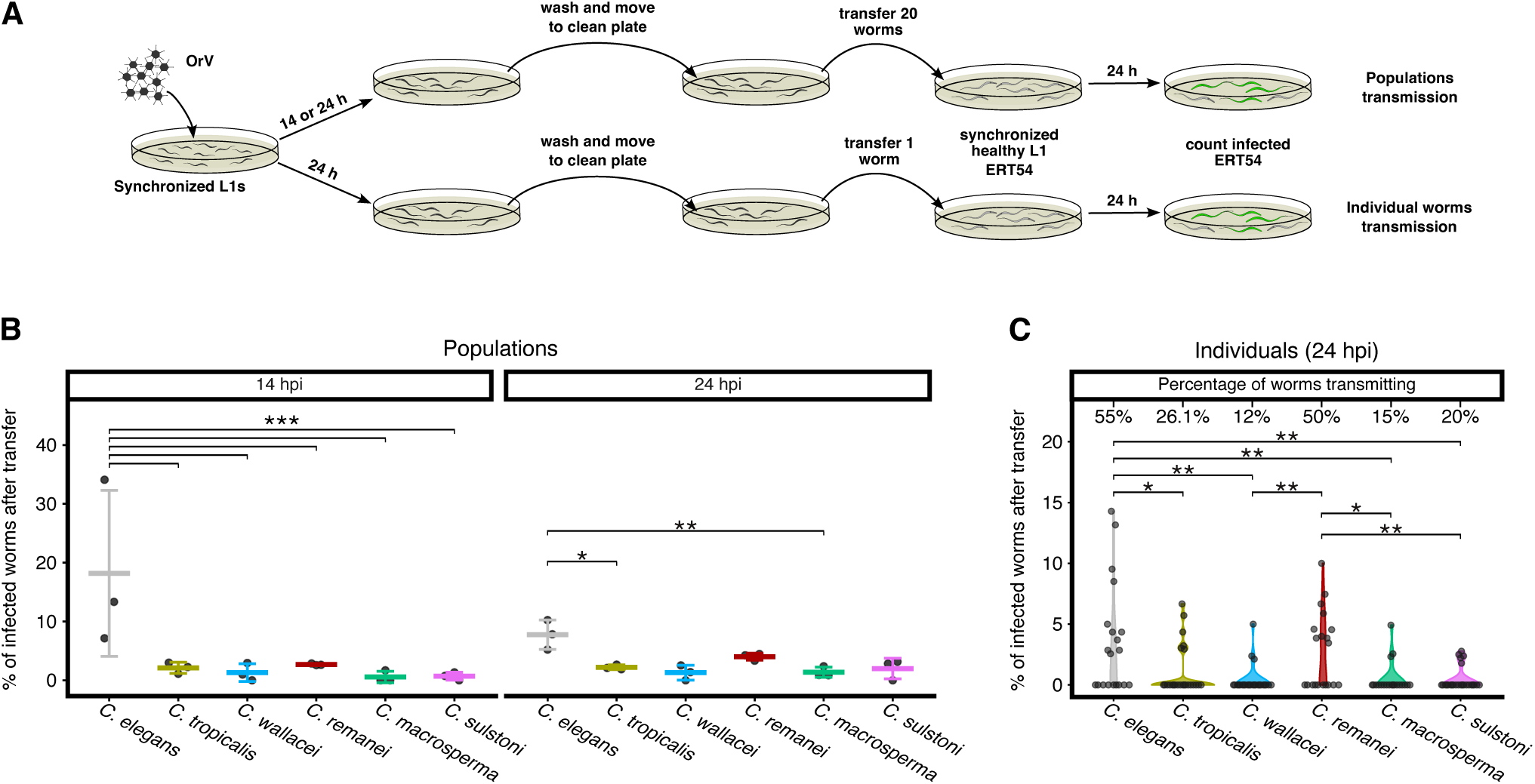
OrV transmission rates are time- and species-specific. (**A**) Experimental design. Synchronized populations of inoculated animals were washed and populations of 20 worms or individual worms were transferred to healthy *C. elegans* ERT54 animals which contain a GPF fluorescent marker of infection. Twenty-four hours after the transfer, the percentage of infected ERT54 animals was determined by counting the number of animals expressing the infection reporter. (**B**) Transmission of OrV from inoculated populations of 20 worms (*n* = 3) of different host species (*C. elegans*, grey; *C. tropicalis*, yellow; *C. wallacei*, blue; *C. remanei*, red; *C. macrosperma*, green; *C. sulstoni*, pink) to populations of 111 ±26 healthy *C. elegans* ERT54 at 14 and 24 hpi. (**C**) Transmission of OrV from individual worms of different host species (*n* = 22 ±3) to populations of 38 ±15 healthy *C. elegans* ERT54 at 24 hpi. The values above each violin plot denotes the percentage of individual worms that successfully transmitted OrV.

Transmission from groups of 20 donors differed significantly among host species (Fig. 4B; χ^2^ = 80.061, 4 d.f., *P* < 0.0001). We also detected a significant interaction between species and the age of the infected donor animals (χ^2^ = 24.886, 6 d.f., *P* = 0.0004), largely driven by a pronounced decrease in transmission between 14 and 24 hpi in *C. elegans* (18.7% *vs* 7.4%, *P* = 0.0003). At 14 hpi, *C. elegans* exhibited the highest transmission probability (18.7%; *C. tropicalis*, 2.2%, *P* < 0.0001; *C. wallacei*, 1.2%, *P* < 0.0001; *C. remanei*, 2.7%, *P* < 0.0001; *C. macrosperma*, 0.6%, *P* < 0.0001; and *C. sulstoni*, 0.8%, *P* < 0.0001). By 24 hpi, however, transmission rates across species were more uniform, with significant differences remaining only between *C. elegans* and *C. wallacei* (1.4%, *P* = 0.0122) and *C. elegans* and *C. macrosperma* (1.4%, *P* = 0.0062).

To assess how many individual worms within a population are able to transmit OrV, we then transferred single donors at 24 hpi into populations of 38 ±15 healthy ERT54 animals (Fig. 4A). The proportion of individuals successfully transmitting infection differed significantly among host species (Fig. 4C; χ^2^ = 13.321, 4 d.f., *P* = 0.0098). These differences reflected two statistically homogeneous groups: a higher-transmission group composed of *C. elegans* (55.0%), *C. tropicalis* (26.1%), *C. remanei* (50.0%), and *C. sulstoni* (20.0%) (*P* ≥ 0.1828 for all comparisons within the group), and a lower-transmission group composed of *C. wallacei* (12.0%) and *C. macrosperma* (15.0%) (*P* ≤ 0.0205). Host species also significantly influenced transmission outcomes (mean fraction of infected recipients; χ^2^ = 37.272, 4 d.f., *P* < 0.0001). *C. elegans* showed significantly higher mean transmission (3.6%) compared to all alternate hosts, except for *C. remanei* (Fig. 4C; *C. tropicalis*, 1.1%, *P* = 0.0286; *C. wallacei*, 0.3%, *P* = 0.0004; *C. remanei*, 2.9%, *P* = 1; *C. macrosperma*, 0.6%, *P* = 0.0031; *C. sulstoni*, 0.5%, *P* = 0.0017).

Together, these results show that OrV transmission, like viral load, is strongly species-specific and exhibits substantial within-species variability. Combining these results with our viral load results, we can notice that RNA accumulation and transmission rate are not well correlated in all species, and that the transmission model might vary in a species-specific way.

### Barrier 6: Virulence and host fitness effects

Although infection rates and viral loads were significantly lower in the alternative hosts, we sought to determine whether the developmental impact of OrV infection was nonetheless species-dependent. To do this, we imaged healthy and inoculated populations from hatching through 72 h and measured body length as a proxy for developmental rate. GLMs were fitted separately for each species.

For the two hermaphroditic species, *C. elegans* (Fig. 5A) showed highly significant effects of infection status and the infection status-by-time interaction (χ^2^ = 86.535, 1 d.f., *P* < 0.0001; χ^2^ = 31.261, 4 d.f., *P* < 0.0001) consistent with previous results (Villena-Giménez et al. 2026). Infected animals were, on average, 5.2% smaller. *C. tropicalis* (Fig. 5B) showed a similar pattern, with both infection status and its interaction with time having strong effects on development (χ^2^ = 57.286, 1 d.f., *P* < 0.0001; χ^2^ = 38.959, 3 d.f., *P* < 0.0001). In this species, infected animals were on average 3.4% smaller.

**Figure 5.**
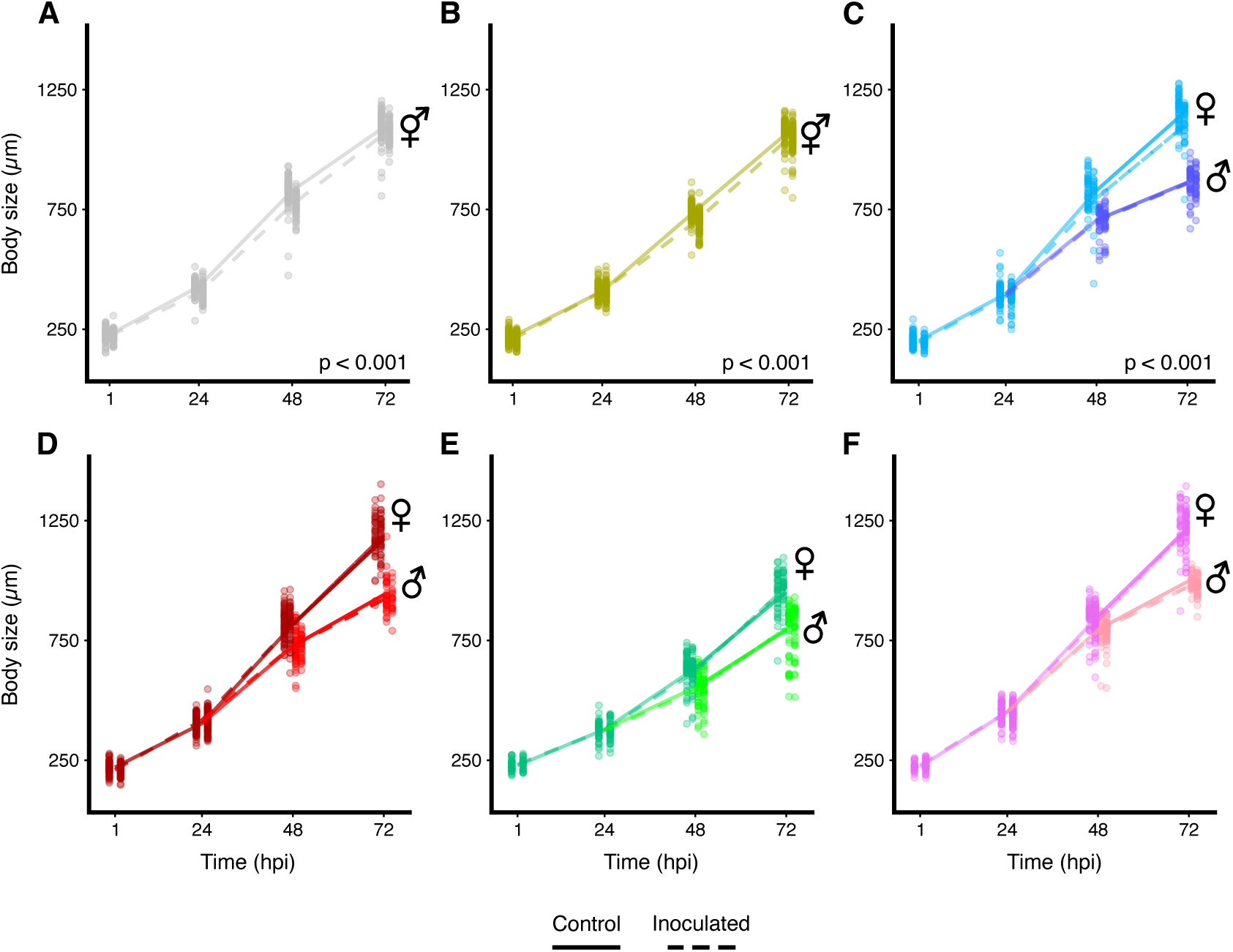
Virulence of OrV is species-specific and can change based on host sex. Virulence measured as the difference in animal body length between inoculated and control populations from hatching to 72 hours. (**A**) *C. elegans* (grey), (**B**) *C. tropicalis* (yellow), (**C**) *C. wallacei* (blue), (**D**) *C. remanei* (red), (**E**) *C. macrosperma* (green), and (**F**) *C. sulstoni* (pink). Mean body length of control populations is represented by full lines and of inoculated populations by dashed lines. Populations of gonochoristic species were separated by sex once it was clearly discernable at 48 and 72 hpi. Males of each gonochoristic species are shown in a different shade color. The sex of each population is displayed next to the corresponding line on the graph.

Because reproductive modes differ among species and sex can only be distinguished reliably from the L4 stage onward, males and females in the gonochoric species were separated at 48 and 72 h. We then tested whether sex contributed to variation in the developmental effects of OrV infection. In *C. wallacei* (Fig. 5C), both infection status (infected animals 2.6% smaller; χ^2^ = 8.125, 1 d.f., *P* = 0.0044) and sex (males 19.6% smaller; χ^2^ = 416.124, 1 d.f., *P* < 0.0001) had significant effects. Infection effects remained constant across these time points (χ^2^ = 0.665, 1 d.f., *P* = 0.4147). Notably, a significant infection status-by-sex interaction (χ^2^ = 7.774, 1 d.f., *P* = 0.0053) indicated that OrV symptoms were more severe in females (4.6% reduction) than in males (0.1%).

Across the other three gonochoric species (*C. remanei*, *C. macrosperma* and *C. sulstoni*; Fig. 5D - F), sex was the primary determinant of body size. Males were consistently smaller, by 13.9% to 16.6%, and also showed slower growth, as indicated by significant sex (χ^2^ ≥ 160.734, 1 d.f., *P* < 0.0001) and sex-by-hpi (χ^2^ ≥ 21.787, 1 d.f., *P* < 0.0001) effects. In none of these species did infection status, alone or in interaction with other factors, have a detectable impact on development (χ^2^ ≤ 3.076, 1 d.f., *P* ≥ 0.0795 in all cases).

Together, these findings indicate that despite low infection rates in non-natural hosts, OrV infection can still exert measurable virulence at the population level.

### Barrier 7: Evolutionary sustainability

After characterizing OrV infection in the alternative host species, we sought to evolve OrV in these six host species. To do this we inoculated five independent lineages of 20 synchronized worms who were left to grow and reproduce until plate starvation at which point OrV was collected to inoculate the next passage. We attempted evolutions with two different versions of the protocol changing the age of the initial animals at inoculation, L1 or L4 (Fig. S3A). OrV viral load in the inoculum was quantified after each passage by RT-qPCR.

When starting passages by inoculating L1s (Fig. S3B), 4/5 OrV lineages evolved in *C. elegans* showed fluctuations in viral load (∼10^4^ to ∼10^6^ OrV RNA2 copies per µL) in passages 1 to 5, and an increase in passage 6 (∼10^7^). The 5^th^ lineage showed a large drop from passage 1 to 2, but the low level of viral load remained stable in a slow decline until passage 6. In the alternative hosts, the viral load of all lineages dropped sharply after passage 1, and fell to levels comparable to negative controls (below ∼10^3^) by passage 3 in *C. macrosperma*, and by passage 4 in *C. tropicalis* and *C. wallacei*. When starting passages by inoculating L4s (Fig. S3C), we observed a similar pattern. All lineages evolved in *C. elegans* showed stable high viral loads (∼10^6^) across the 3 passages performed with this second protocol. All lineages evolved in the alternative hosts once again reached low viral load levels (∼10^3^ or below), this time already by passage 2 and continued declining. Integrating these results with our previous characterization, we conclude that the infection and transmission rates of the alternative host species are not high enough to sustain OrV passaging in these populations using our protocol.

Collectively, mapping outcomes to the barrier framework reveals species-specific blockages at entry/establishment, replication rate/timing, genome stoichiometry, egress, and transmission —with concomitant fitness impacts and failure to sustain evolution in non-natural hosts— thereby explaining why OrV persists and adapts readily in *C. elegans* but not in most alternative *Caenorhabditis* hosts evaluated.

## Discussion

Viral emergence depends on how well a virus’ life cycle aligns with host biology. By dissecting OrV infection across six *Caenorhabditis* species, we show that each host imposes distinct barriers that constrain establishment, transmission and adaptation, which we can interpret through the canonical spillover framework (Plowright et al. 2017). Susceptibility varied widely: some hosts permitted entry but rapidly cleared infection (*e.g*., *C. remanei*), whereas others limited infection to rare individuals (*e.g*., *C. sulstoni* and *C. wallacei*). Frequent RNA1-only cells in restrictive hosts indicate early-cycle bottlenecks in which RNA2 replication fails, blocking virion assembly, precisely the type of molecular mismatch predicted to restrict host range unless viruses surmount rate-limiting steps in replication and packaging in new hosts.

Transmission assays confirmed that competence is multicomponent rather than a simple function of viral load: *C. elegans* remained the most competent transmitter overall; *C. remanei* transmitted surprisingly well given its low loads owing to efficient egress; and species such as *C. tropicalis* and *C. macrosperma* reached high loads yet released few virions. These contrasting host competence phenotypes mirror patterns in wildlife systems where susceptibility, replication and shedding are only weakly correlated (Kilpatrick et al. 2006; Martin et al. 2019). Infection also carried measurable developmental costs in *C. elegans* and *C. tropicalis*, including sex-specific effects in *C. wallacei*, underscoring how demographic structure and mating system shape the ecological impact of infection (Masri et al. 2013; Kailing et al. 2023). Serial passage revealed that most alternative hosts function as evolutionary sinks: OrV rarely persisted beyond a few passages because prevalence, replication, and transmission were insufficient, whereas *C. elegans* consistently maintained high loads and stable persistence, reinforcing its role as OrV’s primary evolutionary reservoir. Taken together, our results show that host range is governed by multi-scale mismatches between viral requirements and host physiology, immunity and life history. Mapping these mismatches to theory provides a mechanistic foundation for understanding how eco-evolutionary forces govern viral emergence (Plowright et al. 2017; Becker et al. 2019) and is consistent with evidence that cross-species transmission is often constrained by genomic and ecological factors, including bottlenecks and fitness valleys (Geoghegan et al. 2016; Geoghegan and Holmes 2018).

OrV replication kinetics in alternative host species diverged markedly from those in *C. elegans*: alternative hosts tended to exhibit prolonged eclipse phases, slower exponential growth and lower peak loads. In our system, viral load and transmission are frequently, but not universally, correlated. We also observed heavy-tailed and, in some cases, bimodal viral load distributions, revealing pronounced within-host heterogeneity. For *C. elegans*, *C. tropicalis* and *C. macrosperma*, rare high-load individuals track a higher fraction of transmitting donors, consistent with a superspreading tail (Pagán et al. 2007; Yang et al. 2021; Kanbayashi et al. 2023). However, despite both *C. tropicalis* and *C. macrosperma* showing the same number of high-load individuals, *C. tropicalis* transmitted more frequently than *C. macrosperma*. For *C. wallacei* and *C. sulstoni* we did not detect high-load outliers (noting limited *n* = 24), and transmission was correspondingly low (Ioannidis et al. 2001; Finke and Conzelmann 2003; Longdon et al. 2015; Inagaki et al. 2016). When present, high load outliers likely act as superspreaders that enable rare spillover opportunities (Lloyd-Smith et al. 2005), while the discordance we document between load and onward transmission across hosts echoes competence heterogeneity in wildlife (Kilpatrick et al. 2006). Completion of the infection cycle further depended on the stoichiometric balance between RNA1 and RNA2. Hosts with skewed RNA2:RNA1 ratios seldom showed lumen-localized virus, implying impaired packaging and limited transmission.

*C. remanei* is an exception: it lacked high-load individuals yet transmitted at levels comparable to *C. elegans*, a pattern explained by our smiFISH data showing a disproportionately high fraction of lumen-localized virus at 14 hpi and relatively balanced RNA2:RNA1 ratios in infected cells. We infer that, in *C. remanei*, OrV completes the cycle efficiently in a narrow time window before immune clearance at ∼24 hpi; part of the subsequent drop in accumulated RNA likely reflects successful packaging and egress rather than failure to replicate. Thus, in *C. remanei*, transmission does not follow a superspreader model but rather a brief, efficient egress model, an alternative competence strategy with implications for emergence (Lloyd-Smith et al. 2005; Kilpatrick et al. 2006; Martin et al. 2016).

Cross-species comparisons of RNA2:RNA1 ratios clarify where hosts block the cycle. In *C. tropicalis*, ratios were not statistically different from *C. elegans* and loads were high, yet we detected no lumen signal; this suggests that entry and/or egress and not replication *per se* are the principal obstacles, consistent with prior reports of high load but lower transmission (Shaw and Kennedy 2025). In *C. wallacei*, the absence of high-load individuals combined with uniformly low RNA2:RNA1 ratios in smiFISH-positive cells argues for compound barriers at entry and replication. Together with *C. remanei*, these cases illustrate that while entry is a common hurdle (Alkan et al. 2024), additional steps (translation, packaging or release) can be host-specific bottlenecks. A useful next step would be to deploy the recombinant OrV system to bypass entry and isolate downstream constraints in each host (Jiang et al. 2014b).

Temporal comparisons further illuminate host control. At early time points (≤ 14 hpi) we detected more smiFISH-positive animals but a higher proportion of RNA1-only cells; by 24 hpi, positivity rates declined in restrictive hosts. Combined with the *C. remanei* pattern (many RNA1-only cells and few positives at 14 hpi; none at 24 hpi), this suggests that infections established with both RNAs are more refractory to clearance, explaining the drop in positive *C. elegans* from 14 to 24 hpi and the complete clearance in *C. remanei* at 24 hpi (Castiglioni et al. 2024a). Several traits appear virus-controlled across hosts, especially the ∼12 hpi first replication peak, whereas others are host-controlled: the frequency of high-load outliers (bimodality), the RNA2:RNA1 balance and egress (Lee et al. 2009; Challenger et al. 2022; Castiglioni et al. 2025). Delayed peaks in *C. tropicalis* and *C. wallacei* and the previously reported sensitivity of peak timing to host developmental stage support a major role for host factors in shaping replication dynamics (Castiglioni et al. 2025, Melero et al. 2025).

Phylogenetic proximity alone does not predict load or virulence in this genus. Although viruses often perform better in closely related hosts and load/virulence can correlate with host phylogeny (Longdon et al. 2011, 2015; Walsh et al. 2023), the high genetic and within-species diversity in *Caenorhabditis*, especially in immunity genes (Ashe et al. 2013), means strain identity can dominate. We observed virulence in *C. tropicalis* and *C. wallacei* (both relatively close to *C. elegans*) despite contrasting loads, no virulence in *C. remanei* (also relatively closely related, low load, brief infection), and no virulence in the more distant *C. macrosperma* and *C. sulstoni* despite higher loads than *C. wallacei*. These patterns indicate that specific host-virus interactions, more than species-level relatedness, determine virulence.

We also found sex-specific effects in *C. wallacei* (females more affected), consistent with broader literature on sex differences in infection outcomes, and paralleling reports that *C. elegans* males show greater resistance to OrV, potentially via higher IPR activation (van der Berg 2006; Masri et al. 2013; van Slujis et al. 2021; Vincent and Dionne 2021).

Finally, our experimental evolution attempts failed in most alternative hosts because each imposes compounding constraints (low prevalence, incomplete cycles or poor egress). Methodology also mattered: our filtration-based passaging selects strictly on viral traits and succeeded in *C. elegans* (see also Castiglioni et al. 2024b), whereas the worm-picking approach used elsewhere may have co-selected host traits increasing permissiveness for infection, helping a minority of lineages persist (Shaw and Kennedy 2022; Shaw and Kennedy 2025).

In sum, the *Caenorhabditis* - OrV system reveals how entry, timing, stoichiometry and egress interact to produce distinct host-competence phenotypes and to determine whether a host is permissive, restrictive or an evolutionary sink. By aligning these outcomes with the spillover framework and with theory on hierarchical barriers, bottlenecks and heterogeneity (Lloyd-Smith et al. 2005; Kilpatrick et al. 2006; Geoghegan et al. 2016; Plowright et al. 2017; Geoghegan & Holmes 2018; Becker et al. 2019), we provide a mechanistic basis for predicting host range and emergence and a roadmap for targeted experiments (*e.g*., entry bypass or host factor perturbations) to pinpoint the step that fails in each host.

## Methods

### Worm strain maintenance

*C. elegans* strains ERT54 and JU2624 were obtained from E.R. Troemel and M.A. Félix, respectively. *C. elegans* N2, *C. tropicalis* JU1428, *C. macrosperma* JU1857, *C. wallacei* JU1873, and *C. sulstoni* SB454 were obtained from the Caenorhabditis Genetics Center (CGC, https://cgc.umn.edu). The genetically diverse *C. remanei* SP8 generated through artificial selection (Chen and Maklakov 2012; Lind 2024) was obtained from M. Lind. For *C. elegans*, the ERT54 strain was used in experiments unless stated otherwise. All strains were maintained under standard conditions (Brenner 1974; Stiernagle 2006) at 20 °C on NGM plates seeded with live *Escherichia coli* OP50. *C. elegans*, *C. tropicalis*, *C. wallacei*, and *C. remanei* belong to the *Elegans* group, while *C. macrosperma* and *C. sulstoni* belong to the *Japonica* group (Stevens et al. 2020).

### OrV stock generation and quantification

OrV stock was generated by inoculating two 6 cm plates of freshly starved JU2624 worms with the JUv1580_vlc (GenBank: PP738529.1 for RNA1 and PP738530.1 for RNA2) OrV strain and expanding them on fifty 9 cm NGM plates until starvation. Plates were washed with M9 buffer (0.22 M KH_2_PO_4_, 0.42 M Na_2_HPO_4_, 0.85 M NaCl, 1 mM MgSO_4_) and pooled into 15 mL tubes. Tubes were vortexed for 30 s, incubated at room temperature for 10 min, and centrifuged at 2,250×g for 2 min. The worm pellet was discarded, and the supernatant was transferred to a new tube, centrifuged twice at 21,000×g for 5 min at 4 °C, and filtered through a 0.22 µm syringe filter. The resulting viral preparation was stored at −80 °C.

Viral RNA was extracted using the Viral RNA Isolation Kit (NZYtech, Lisbon, Portugal) following the manufacturer’s instructions. OrV titers were quantified by standard curve RT-qPCR using the qPCRBIO SyGreen 1-Step Go Hi-ROX kit (PCR Biosystems Ltd., London, UK) on an ABI StepOnePlus Real-Time PCR System (Applied Biosystems, Foster City CA, USA), with primers targeting a region of OrV RNA2 (5’ACGAAGCAGTAGCCGTTAAG3’ and 5’GAGAACATCCTTCTCTGCGG3’).

The standard curve consisted of six serial dilutions ranging 1.25×10^9^ - 1.25×10^4^ copies of OrV RNA2/µL, generated from an *in vitro* transcribed RNA2 fragment. cDNA for *in vitro* transcription was synthesized using AccuScript High Fidelity Reverse Transcriptase (Agilent, Santa Clara CA, USA) and a reverse primer binding the 3’ end of RNA2 (5’ATAGCCGGGTATGGATAGCG3’). Approximately 1,000 bp from the 3’ region were amplified using a forward primer containing a 20 bp T7 promoter sequence (5’TAATACGACTCACTATAGGCCTGTCAGAGTTGAGAACA3’) and DreamTaq DNA Polymerase (Thermo Fisher Scientific, Waltham MA, USA). The PCR product was gel-purified using the MSB Spin PCRapace kit (Invitek Molecular GmbH, Berlin, Germany) and transcribed *in vitro* using T7 polymerase (Merck & Co., Rahway NJ, USA).

### Worm synchronization and inoculation

Synchronized worm populations were obtained by washing plates with M9 buffer to remove adults and larvae while leaving eggs behind. Plates were incubated at 20 °C for 1 h, and newly hatched larvae were collected with M9 and transferred to new plates. Larvae were inoculated by applying OrV stock containing a total of 2.8×10^9^ copies of RNA2 onto the *E. coli* OP50 lawns. Control plates received an equivalent volume of M9 buffer.

### Total RNA extraction and RT-qPCR

Populations of ∼500 inoculated or control worms were collected using PBS with 0.05% Tween-20. Samples were centrifuged at 400×g for 2 min, washed three times, flash-frozen in liquid N_2_, and stored at −80 °C overnight.

For lysis, 500 µL of TRIzol (Thermo Fisher Scientific) or EasyBlue (iNtRON Biotechnology, Kirkland WA, USA) were added, followed by five freeze-thaw cycles between liquid nitrogen and 37 °C. Samples were vortexed five times for 30 s with 30 s rests. Chloroform (100 µL) was added, samples were mixed for 15 s, incubated 3 min at room temperature, and centrifuged at 16,000×g for 15 min at 4 °C. The aqueous phase was transferred to new tubes, mixed with an equal volume of ethanol, and RNA was purified using the RNA Clean & Concentrator-5 Kit (Zymo Research, Orange CA, USA).

RNA concentration was measured with a NanoDrop OneC spectrophotometer (Thermo Fisher Scientific), and samples were diluted to 10 ng/µL. RT-qPCR was performed using 10 ng of total RNA as described above.

### Single-worm lysis and RT-qPCR

Single-worm lysis was adapted from Ly et al. (2015). Synchronized infected and control worms were washed three times with PBS + 0.05% Tween-20 and transferred to NGM plates without *E. coli* OP50. Individual worms were picked into 0.2 mL tubes containing 5 µL of lysis buffer (5 mM Tris pH 8.0, 0.25 mM EDTA, 0.5% Triton X-100, 0.5% Tween-20, 0.1% proteinase K). Lysis was performed in a thermocycler at 65 °C for 10 min and 85 °C for 1 min. Lysates were stored immediately at −80 °C. RT-qPCR was performed using 1 µL of lysate as template.

### Transmission assays

Synchronized, inoculated worms were washed three times with PBS + 0.05% Tween-20 and once with M9, then transferred to fresh plates. Worms were picked onto plates containing healthy *C. elegans* ERT54 worms (Bakowski et al. 2014). For *C. elegans*, the N2 strain was used as donor to avoid confusion with the recipient ERT54 animals because the two strains do not show differences in viral load (Fig. S4; Kruskal-Wallis test, χ^2^ = 0.365, 1 d.f., *P* = 0.5457). After 24 h, ERT54 worms were examined for infection by GFP fluorescence using a MZ10F stereomicroscope (10×/23B objective; mCherry M10F/MZ FLII and GFP3 MZ10F filters) (Leica Microsystems GmbH, Wetzlar, Germany). The percentage of GFP-positive ERT54 worms was calculated.

### smiFISH

The smiFISH protocol was adapted from Tsanov et al. (2016) and Parker et al. (2021). Populations of ∼700 inoculated and control worms were washed using the same procedure as for RNA extraction, except with one additional wash and centrifugation at 400×g for 1 min. After the final wash, samples were fixed with 400 µL Bouin’s fixative, 400 µL 100% methanol, and 1 µL β-mercaptoethanol, mixed for 30 min at room temperature on a rotary shaker, flash-frozen, and stored at −80 °C overnight or longer.

Samples were then thawed at 4 °C for 30 min and washed four times with 1 mL borate-Triton buffer (20 mM H_3_BO_3_, 10 mM NaOH, 0.5% Triton X-100) and five times with the same buffer containing 2% β-mercaptoethanol. The last three washes included 1 h incubations at room temperature.

Samples were incubated for 5 min in wash buffer A with formamide (20% Stellaris Wash Buffer A, 20% formamide, 60% water), centrifuged, and resuspended.

Twenty-two probes targeting OrV RNA1 and 18 targeting RNA2 were designed using Oligostan (Tsanov et al. 2016). Probe sequences can be found elsewhere (Castiglioni et al. 2024a, 2024b, 2025; Melero et al. 2025). Probes (0.83 µM) were annealed with a FLAP-label (CAL Fluor 610 or Quasar 670) at 85 °C for 3 min, 65 °C for 3 min, and 25 °C for 5 min. Hybridization was performed overnight at 37 °C in the dark.

The next day, samples were washed with wash buffer A, incubated 30 min at 37 °C, stained with 25 ng DAPI, washed with Stellaris Wash Buffer B, resuspended in 70 µL of Buffer B with 0.1 ng DAPI, and. mounted with N-propyl gallate. For each replicate, 56 ±4 animals were randomly selected and imaged using a DMi8 microscope with a DFC9000 GTC sCMOS camera (Leica Microsystems GmbH). Images were processed in Fiji (Schindelin et al. 2012).

Animals were counted as infected if at least one of the OrV RNAs was detected. Infected animals were analyzed by measuring the distance from the pharyngeal-intestinal junction to the start and end of the infected region. Relative infected area was calculated as the infected length divided by the total intestinal length. The median fluorescence intensity of infected cells was measured for both RNA1 and RNA2. Background values were obtained from control worms. Median fluorescence intensities were background-corrected and normalized by the number of probes used for each RNA, and RNA2:RNA1 ratios were calculated.

### Growth images and measurements

Populations of 80 ±25 synchronized infected and control worms were transferred to NGM plates with *E. coli* OP50. Plates were imaged at designated time points using a MZ10F stereomicroscope and Flexacam C3 camera (Leica Microsystems GmbH). Worm length was measured in Fiji by tracing the midline from the anterior tip of the head to the posterior end of the body, excluding the tail.

### Experimental evolution

The first passage was started with populations of 20 synchronized L1 or L4 worms which were inoculated with 2.8×10^9^ OrV RNA2 copies in NGM plates seeded with 200 µL of *E. coli* OP50. The populations were allowed to grow and reproduce until plate starvation at which point worms and virus from each plate were collected with M9 buffer. Each sample was vortexed 3 times for 30 s with 30 s rests and left to incubate at room temperature for 10 min. Samples were then centrifuged for 2 min at 400×g and 4 °C. The supernatant was then centrifuged for 5 min at 12,000×g and 4 °C, filtered through a 0.22 µm syringe filter, and stored at −80 °C. For each subsequent passage, worms and plates were prepared as described for the first passage, however, inoculation was done with 100 µL of OrV filtrate from the previous passage. The OrV filtrate from each passage was quantified by standard curve RT-qPCR as described for OrV stock quantification above.

### Statistical analysis

Statistical analyses were performed in R version 4.5.2 using RStudio Desktop version 2026.01.0+ (Posit, Boston, MA, USA) or in SPSS version 30.0 (IBM, Armonk, NY, USA).

Log_10_-transformed viral load data from population experiments were analyzed using generalized linear models (GLMs) with a Normal distribution and identity-link function. Nematode species was included as a fixed factor nested within its corresponding taxonomic group (*Elegans* or *Japonica*). Time post-infection (hpi) was incorporated as an orthogonal random factor, and the species-by-time interaction was also evaluated. For single-animal viral load estimates, hpi was not included as a factor. To characterize the heavy-tailed viral load distributions, we fitted two alternative models: (*i*) a unimodal Lognormal distribution with two parameters, and (*ii*) a mixture of two Lognormal distributions with weights *p*_1_ and 1 – *p*_1_ and four additional parameters. Model fitting was performed using the Levenberg-Marquardt algorithm.

Transmission data from population experiments were analyzed using GLMs with a Binomial distribution and logit-link function. Species was treated as a fixed factor nested within taxonomic group, and host age at inoculation (14 or 24 h) was included as an orthogonal random factor. The interaction between species and age was also tested. For single-animal transmission assays, age was not included as a factor.

Infectivity data from smiFISH experiments at 14 or 24 hpi were analyzed using GLMs with a Binomial distribution and logit-link function, with species treated as a fixed factor nested within taxonomic group.

The effects of nematode species on the area and relative position of infected cells along the body axis at 24 hpi were analyzed using GLMs with a Normal distribution and identity-link function, with species nested within taxonomic group as a fixed factor. Comparisons between *C. elegans* and *C. remanei* at 14 hpi were conducted using Mann-Whitney *U* tests.

Counts of infected cells per animal at 14 and 24 hpi were analyzed using GLMs with a Poisson distribution and log-link function, with species nested within taxonomic group as a fixed factor.

The presence of virus in distinct anatomical locations (cells, lumen or both) was analyzed using GLMs with a Multinomial distribution and an accumulated logit-link function, with species treated as a fixed factor nested within taxonomic group.

Ratios of normalized RNA2 to RNA1 fluorescence at 24 hpi were analyzed using GLMs with a Normal distribution and identity-link function, with species nested within taxonomic group. Comparisons between *C. elegans* and *C. remanei* at 14 hpi were performed using Mann-Whitney *U* tests.

Finally, developmental size data were analyzed by fitting GLMs with a Normal distribution and identity-link function, including infection status and hpi as orthogonal factors. Biological replicate was nested within the interaction between the two orthogonal factors. For gonochoristic species (*C. wallacei*, *C. remanei*, *C. macrosperma*, and *C. sulstoni*), gender was included as an additional orthogonal factor.

In all analyses, *post hoc* pairwise comparisons were conducted using the Holm-Bonferroni method. Additional statistical tests are described where relevant in the text.

## Acknowledgements

We thank Francisca de la Iglesia for excellent technical support and the members of the EvolSysVir lab for valuable comments and fruitful discussions. We also thank Wormbase and the Caenorhabditis Genetics Center. This work was supported by grants PID2022-136912NB-I00 funded by MCIU/AEI/10.13039/501100011033 and by “ERDF a way of making Europe”, and CIPROM/2022/59 funded by Generalitat Valenciana to S.F.E. D.H. was supported by grant PREP2022-000699 funded by MCIU/AEI/10.13039/501100011033 and by “ESF investing in your future”. J.C.M-S. was supported by grant ACIF/2021/296 from Generalitat Valenciana. V.G.C. was supported by grant MSCA 2024-PF-01-101207897 funded by Horizon Europe.

## Author contributions

D.H.: Writing-original draft, conceptualization, investigation, writing-review and editing, methodology, data curation, formal analysis, resources, and visualization. J.C.M-S.: Conceptualization, writing-review and editing, formal analysis, and visualization. A.V-G.: Conceptualization, writing-review and editing, and methodology. V.G.C.: Writing-original draft, conceptualization, writing-review and editing, methodology, supervision, validation. S.F.E.: Writing-original draft, conceptualization, writing-review and editing, formal analysis, funding acquisition, data curation, validation, supervision, and project administration.

## Data accessibility

All raw data generated in this study can be accessed at Zenodo (https://doi.org/10.5281/zenodo.19473000).

---

**Supplementary Figure S1.** Distribution of viral loads of individual animals and corresponding fitted probability density functions. OrV viral load of individual animals (*n* = 24 for each species). (A) *C. elegans* ERT54 (grey), (B) *C. tropicalis* JU1428 (yellow), (C) *C. wallacei* JU1873 (blue), (D) *C. remanei* SP8 (red), (E) *C. macrosperma* JU1857 (green), and (F) *C. sulstoni* SB454 (pink) at 24 hpi. Viral load was measured as the number of OrV RNA2 copies in 1/5 of the single worm lysate. Horizontal and vertical grey shaded bands correspond to one standard deviation around the control-group mean viral load. First row: individual viral load measurements are shown as dots, together with a violin plot that highlights the regions of highest data density within each strain. Second row: fitted probability density function with the original viral load variable, using either a Lognormal or a bimodal Lognormal distribution depending on the strain (Table 1). Third row: the same fitted distributions shown in logarithmic space, where they correspond to normal or bimodal normal distributions for log_10_-viral load. In both cases, the displayed curves are derived from the same fitted parameters.

**Supplementary Figure S2.** Representative images of distinct infection phenotypes observed by smiFISH fluorescence microscopy. *C. elegans* worms at 14 or 24 hpi showing (A) infected animals with OrV localized in a cell, in the intestinal lumen, or in both, and (B) infected animals with 1, 2, 3, or 4 infected cells. The brightness of each image was adjusted separately.

**Supplementary Figure S3.** Experimental evolution of OrV in *C. elegans* and alternative hosts. (A) Experimental design. Five independent lineages of OrV were passaged in each host. Synchronized populations of 20 animals were inoculated with 2.8×10^9^ copies of OrV RNA2 to start the first passage. Once starved, plates were washed and evolved virus was filtered out. To inoculate each subsequent passage, 100 µL of virus filtrate from the previous passage was used. There were two experimental regimes that differed in the age of the worms which were inoculated to start a passage, either L1 or L4. (B) OrV viral load, measured as number of RNA2 copies per µL, after each passage of experimental evolution starting with L1-inoculated animals. (C) OrV viral load, after each passage of experimental evolution starting with L4-inoculated animals. *C. elegans* ERT54 (grey), *C. tropicalis* JU1428 (yellow), *C. wallacei* JU1873 (blue), *C. remanei* SP8 (red), *C. macrosperma* JU1857 (green), and *C. sulstoni* SB454 (pink).

**Supplementary Figure S4.** Individual worm viral load of *C. elegans* strains ERT54 and N2. OrV viral load of individual animals (*n* = 24 for each strain) at 24 hpi. Viral load was measured as the number of OrV RNA2 copies in 1/5 of the single worm lysate. Horizontal grey line represents the technical detection limit (mean + 2 SD of negative controls). The viral loads of the two strains do not differ significantly (Kruskal-Wallis test, χ^2^ = 0.365, 1 d.f., *P* = 0.5457).

