## Supplementary figures and images for "Species-specific barriers constrict Orsay virus host range across the *Caenorhabditis* genus"

### Supplementary Figures

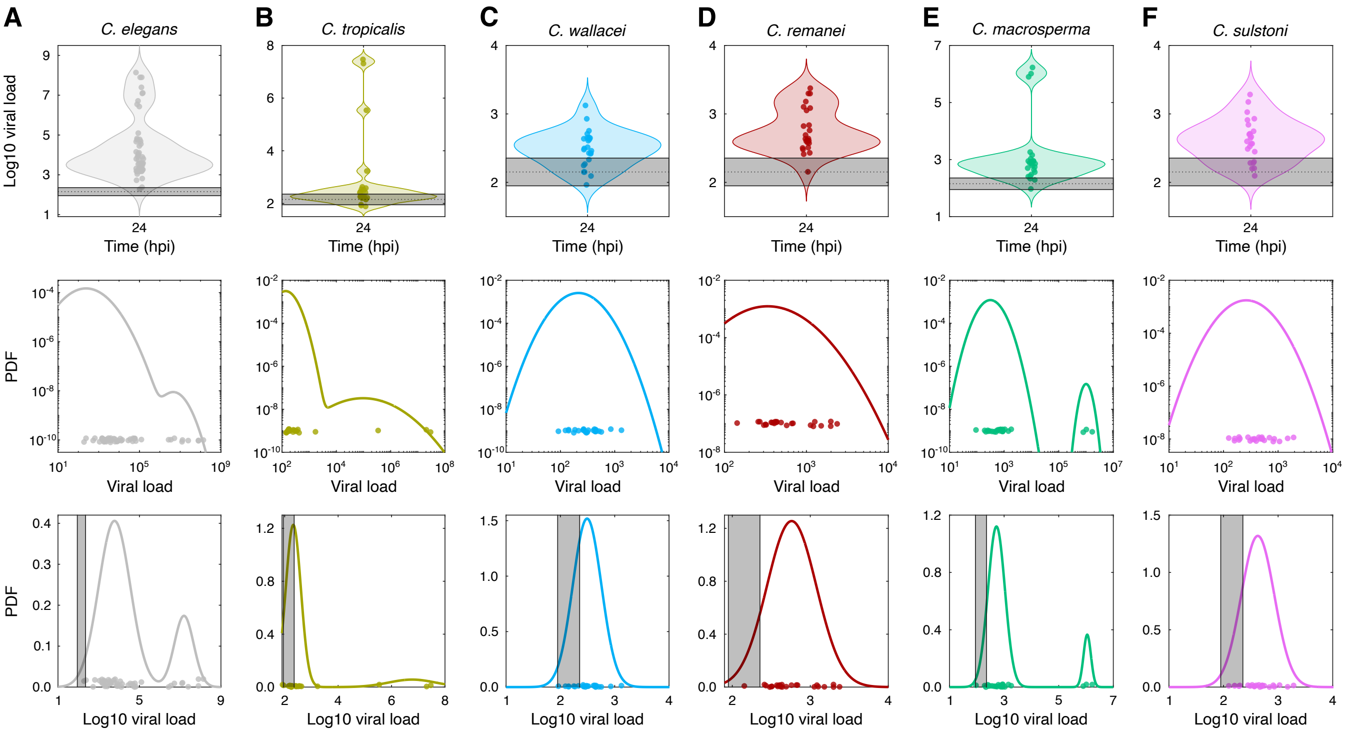

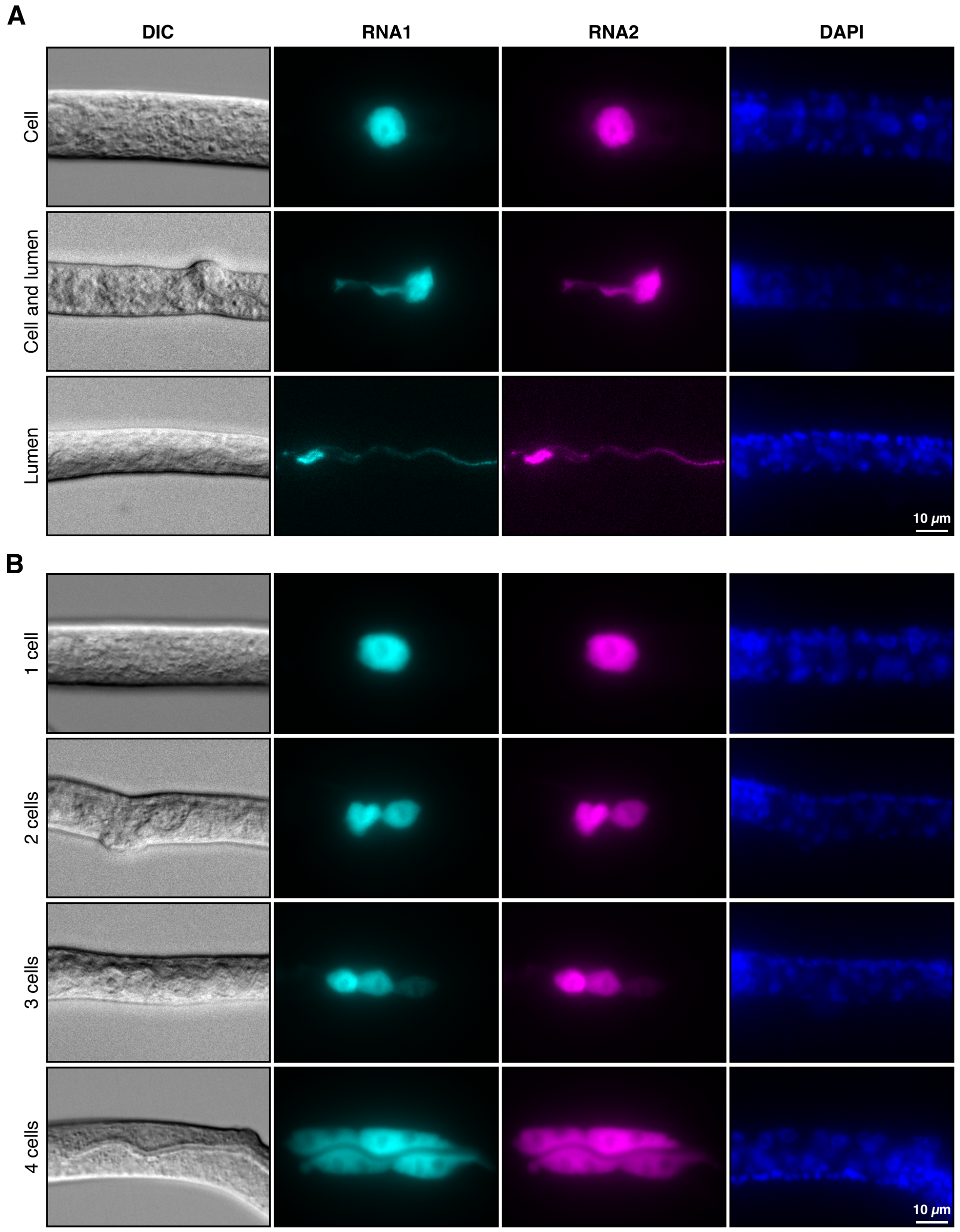

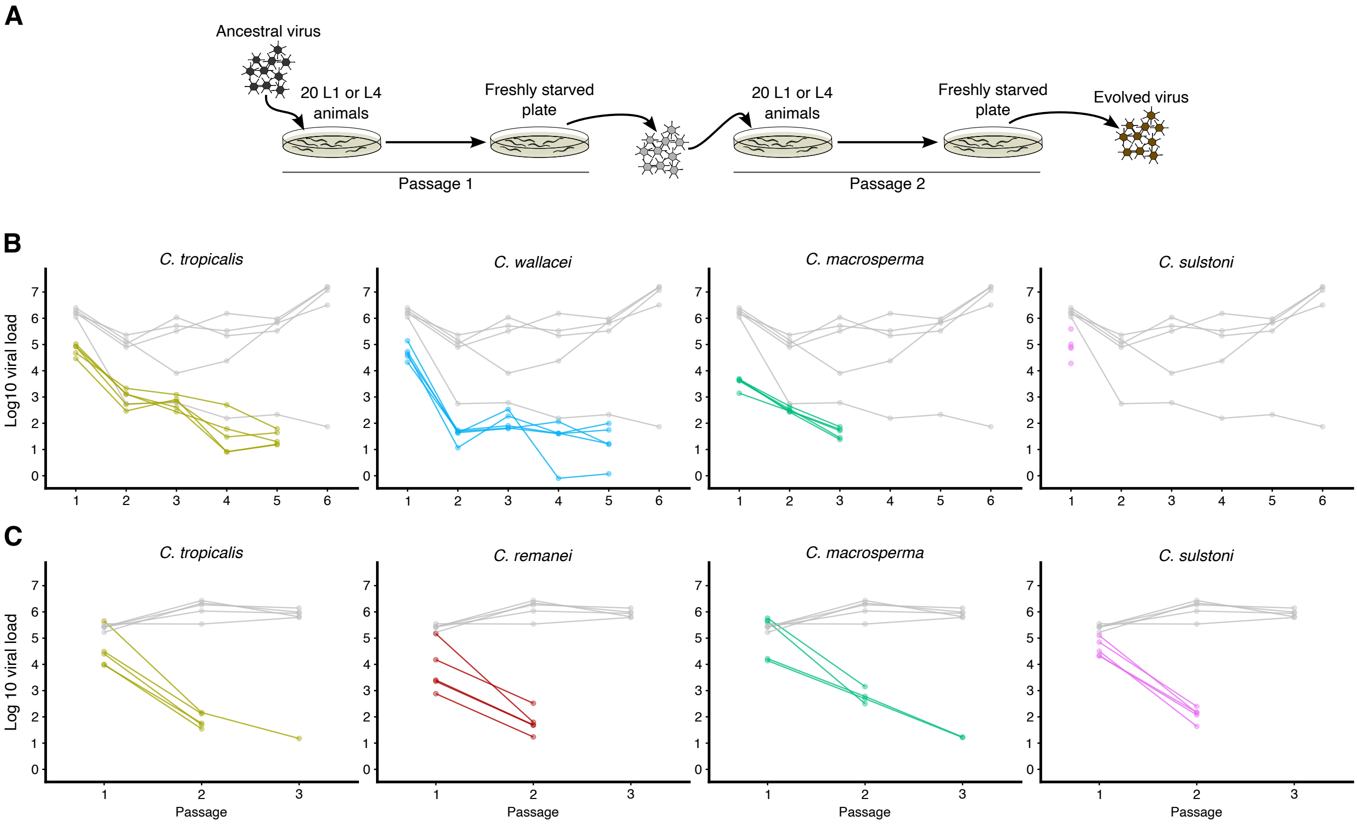

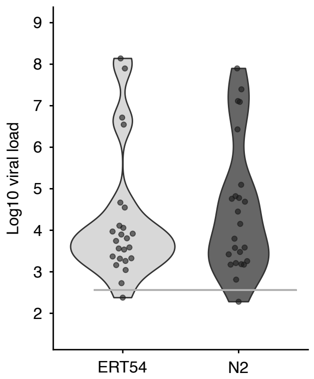
